# Mammals adjust diel activity across gradients of urbanization

**DOI:** 10.1101/2021.09.24.461702

**Authors:** Travis Gallo, Mason Fidino, Brian Gerber, Adam A. Ahlers, Julia L. Angstmann, Max Amaya, Amy L. Concilio, David Drake, Danielle Gray, Elizabeth W. Lehrer, Maureen H. Murray, Travis J. Ryan, Colleen Cassady St. Clair, Carmen M. Salsbury, Heather A. Sander, Theodore Stankowich, Jacque Williamson, J. Amy Belaire, Kelly Simon, Seth B. Magle

**Affiliations:** Environmental Science and Policy, College of Science, George Mason University, Fairfax, VA 22030 USA; Urban Wildlife Institute, Conservation and Science Department, Lincoln Park Zoo, Chicago, IL 60614 USA; Department of Natural Resource Science, The University of Rhode Island, Kingston, RI 02881, USA; Department of Horticulture and Natural Resources, Kansas State University, Manhattan, KS 66502 USA; Department of Biological Sciences and Center for Urban Ecology and Sustainability, Butler University, Indianapolis, IN 46208 USA; Department of Biological Sciences, California State University Long Beach, Long Beach, CA 90840 USA; Department of Environmental Science and Policy, St. Edward’s University, Austin, TX 78704 USA; Austin Parks and Recreation, City of Austin, TX 78704 USA; Department of Forest and Wildlife Ecology, University of Wisconsin-Madison, Madison, WI 53706, USA; Department of Biological Sciences, University of Alberta, Edmonton, Canada; Department of Geographical and Sustainability Sciences, University of Iowa, Iowa City, IA 52242 USA; Department of Education & Conservation, Brandywine Zoo, Wilmington, Delaware, 19802 USA; The Nature Conservancy in Texas, San Antonio, Texas 78215 USA; Texas Parks and Wildlife, Austin, Texas 78774 USA

**Keywords:** behavior, human disturbance, nocturnality, temporal partitioning, urban wildlife

## Abstract

Time is a fundamental component of ecological processes. How animal behavior changes over time has been explored through well-known ecological theories like niche partitioning and predator-prey dynamics. Yet, changes in animal behavior within the shorter 24-hour light-dark cycle have largely gone unstudied. Understanding if an animal can adjust their temporal activity to mitigate or adapt to environmental change has become a recent topic of discussion and is important for effective wildlife management and conservation. While spatial habitat is a fundamental consideration in wildlife management and conservation, temporal habitat is often ignored. We formulated a temporal resource selection model to quantify the diel behavior of eight mammal species across ten U.S. cities. We found high variability in diel activity patterns within and among species and species-specific correlations between diel activity and human population density, impervious land cover, available greenspace, vegetation cover, and mean daily temperature. We also found that some species may modulate temporal behaviors to manage both natural and anthropogenic risks. Our results highlight the complexity with which temporal activity patterns interact with local environmental characteristics, and suggest that urban mammals may use time along the 24-hour cycle to reduce risk, adapt, and therefore persist in human-dominated ecosystems.

## Introduction

Time is a fundamental axis that shapes ecological systems. Regarding animal behavior, time and space are linked in that the spatial characteristics of an animal’s local environment influences its temporal behavior (Kronfeld-Schor and Dayan, 2003). For example, some species make seasonal changes in diel (24-hour period) activity to be most active during optimal temperatures in their local environment (Maloney et al., 2005), and other species temporally partition themselves from heterospecific competition or aggression (Kronfeld-Schor and Dayan, 2003; van der Vinne et al., 2019). While temporal behavior has yet to become a major focus in animal ecology (Gaston, 2019; Kronfeld-Schor and Dayan, 2003), how animals use time as an ecological resource has inspired well-known ecological phenomenon like niche partitioning (Schoener, 1974) and predator-prey dynamics (Tambling et al., 2015). From an applied perspective, understanding if an animal can make temporal adjustments to mitigate or adapt to local environmental change remains a topic of discussion (Wolkovich et al., 2014), and is important for effective wildlife management and conservation (Levy et al., 2019).

Species that persist in human-dominated environments, like cities, require some degree of human avoidance to safely navigate these complex landscapes (Gehrt et al., 2009; Murray and St. Clair, 2015; Riley et al., 2003). In urban ecosystems, few habitat patches remain for animals to seek spatial refuge when confronted with human disturbance and/or negative interactions with other species. In these cases, temporally partitioning from these potentially dangerous interactions might be an alternative strategy. A recent global meta-analysis suggests that mammals become more nocturnal in areas with greater human disturbance (Gaynor et al., 2018). However, only 7.8% (n = 11) of these studies assessed changes in nocturnal activity in urban areas, and all explored these changes categorically between urban and non-urban areas. Binary urban and rural categorizations generally fail to capture variation in urban development and cannot generate generalizable results that correlate to other cities (McDonnell and Pickett, 1990). Additionally, cities are unique and differ in size, land use, growth patterns, and human culture (Pacione, 2009). Variation in both spatial and temporal characteristics within and among cities could have differing effects on animal behavior. Thus, key questions remain regarding the way in which animal diel activity varies across gradients of urbanization and among differing cities. For example, the magnitude of change in diel activity patterns may be larger for more densely urbanized cities or may depend on regional variation in day and night-time temperatures. Multi-city investigations that include variation in urban intensity and regional climate can elucidate such patterns.

Gaynor et al. (2018) found that most studies in urban environments also focused on carnivore species, highlighting a gap in our understanding regarding changes in diel activity across taxa. For example, carnivores likely avoid humans in both space and time because of inimical human interactions (Clinchy et al., 2016; Kitchen et al., 2000). This may not be the case for mammals that do not regularly come in conflict with humans or do not evoke such visceral reactions by humans. Additionally, some species may be constrained by their morphology (e.g., number and type of cones and rods in their eyes) or may otherwise lack the ability to be active in alternative lighting. To fully understand the variability of activity patterns and assess temporal adjustments in response to urban development, a comprehensive examination of the larger suite of urban mammals and across multiple urban environments is required.

While spatial habitat is a fundamental consideration in wildlife management and conservation, temporal habitat is often ignored (Gaston, 2019). Here, we link spatial landscape characteristics with the diel activity patterns of eight terrestrial mammals using remote cameras deployed across ten U.S. cities. Our objectives were to 1) determine which species change their diel activity across gradients of urbanization and identify what characteristics of the urban environments have the strongest association with changes in diel activity and 2) assess whether urbanization influences nocturnal behavior and identify what characteristics of urban environments have the strongest influence on changes in nocturnal behavior.

We found high variability in diel activity patterns within and among species and species-specific correlations between diel activity and human population density, impervious land cover, available greenspace, vegetation cover, and mean daily temperature. Our results indicate that in high-risk environments, such as cities, animals may reduce risk by modulating their temporal habitat use. Our study identifies a potential mechanism by which urban wildlife species may adapt to human-dominated environments, and provides critical insight into activity patterns of urban wildlife that will prove useful for managing these species in cities

## Results

To quantify changes in mammal diel activity in response to urbanization, we used camera detection data for eight common urban mammal species: bobcat (*Lynx rufus*), coyote (*Canis latrans*), red fox (*Vulpus vulpus*), raccoon (*Procyon lotor*), striped skunk (*Mephitis mephitis*), eastern cottontail (*Sylvilagus floridanus*), Virginia opossum (*Didelphis virginiana*), and white-tailed deer (*Odocoileus virginianus*). Cameras were deployed in a systematic fashion across ten U.S. metropolitan areas as part of the Urban Wildlife Information Network: Austin, TX, Chicago, IL, Denver, CO, Fort Collins, CO, Indianapolis, IN, Iowa City, IA, Orange County, CA, Madison, WI, Manhattan, KS, and Wilmington, DE (and Fidino et al., 2021 for details; see Magle et al., 2019).

Across 41,594 trap nights (Table S1), we captured 79,659 total unique detection events. Total detections per species ranged from 102-34,931, and each species was detected in 5-10 cities at an average proportion of 0.16-0.77 sites per city (Table 1). Bobcat occurred at the lowest number of cities and proportion of sites, while raccoon occurred in all 10 cities and at the greatest proportion of sites (Table 1, see Table S2 for the proportion of sites in each city). The number of detections captured throughout the 24-hour diel period varied among species (Table 1).

**Table 1.** The total number of detections for each species, number of cities each species was detected in, mean proportion of sites each species was detected at per city, and total number of detections in each time category for eight urban mammal species across ten U.S. metropolitan areas between January 2017 and December 2018.

| Species | Total<br>detections | No. of<br>cities<br>species<br>detected | Mean<br>proportion of<br>sites species<br>detected per<br>city | No. of 'day'<br>detections | No. of<br>'dawn'<br>detections | No. of<br>'dusk'<br>detections | No. of<br>'night'<br>detections | No. of<br>'darkest<br>night'<br>detections |
| --- | --- | --- | --- | --- | --- | --- | --- | --- |
| Bobcat | 102 | 5 | 0.16 | 29 | 1 | 9 | 45 | 18 |
| Coyote | 2732 | 9 | 0.63 | 671 | 98 | 256 | 1318 | 389 |
| Eastern cottontail | 16102 | 10 | 0.61 | 3984 | 619 | 1097 | 8317 | 2085 |
| Raccoon | 34931 | 10 | 0.77 | 2638 | 642 | 3767 | 21723 | 6161 |
| Red fox | 1570 | 8 | 0.51 | 441 | 35 | 152 | 744 | 198 |
| Striped skunk | 990 | 10 | 0.24 | 89 | 24 | 98 | 584 | 195 |
| Virginia opossum | 8357 | 8 | 0.7 | 407 | 116 | 1027 | 5087 | 1720 |
| White-tailed deer | 14875 | 10 | 0.56 | 7965 | 658 | 816 | 4299 | 1137 |

### Modeling diel activity

We formulated a hierarchical multinomial model to quantify the diel behavior of each species and assess the effects that available greenspace, vegetation cover, impervious land cover, human population density, and daily temperature had on diel behavior of each species. Our approach operates similar to resource selection functions in which resources are selected in space. However, substituting time for space allowed us to quantify changes in diel activity across gradients of environmental change. This temporal resource selection model allowed us to estimate temporal ‘selection’ and the probability of ‘use’ in each time category. Coefficient estimates are estimates of selection for a particular time category relative to the available time in the respective category and the difference from the reference time category (‘day’). Exponentiated coefficient estimates greater than one indicates selection and less than one indicates avoidance, relative to the day reference category. Using the softmax function (Kruschke, 2011), we also estimated the influence that each predictor variable had on the probability of activity in each time category, including the ‘day’ category.

### Among city variation in diel activity patterns

We found that most species, on average, had a higher probability of being nocturnal (active at night or during the darkest portions of night) with the exception of bobcat and white-tailed deer (Fig. 1 and 2). Most species showed variation in diel activity among cities (e.g., bobcat; Fig. 1), and some species (e.g., eastern cottontail, coyote, red fox, and bobcat) exhibited profound variation in diel activity across individual sampling sites (Fig. 2). For example, the predicted probability of nocturnal behavior for eastern cottontail at each sampled site ranged from 0.15 – 0.69 (see Table S3 for a full set of ranges for each species and each time category).

**Figure 1.**
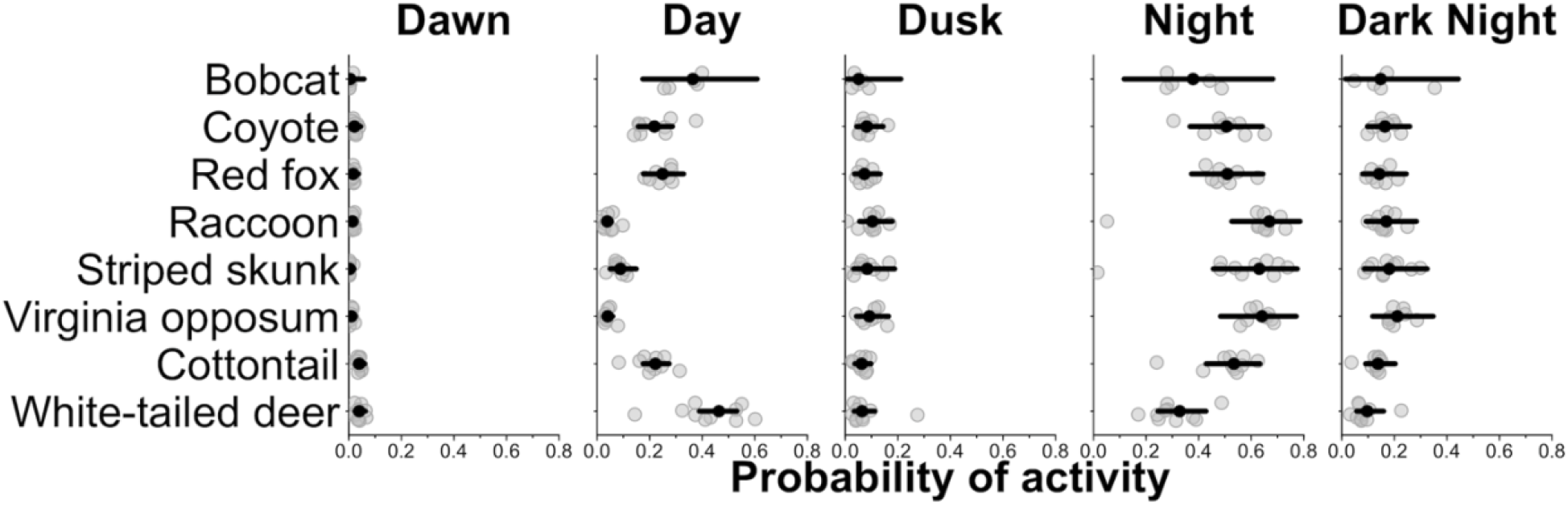
City-specific probability of activity for each species. Grey points are city specific estimates of the average probability of activity in each time category. The black point indicates the average probability of activity among cities and the horizontal lines are 95% credible interval for the average probability estimates among cities. Wider credible intervals indicate more variation among cities.

**Figure 2.**
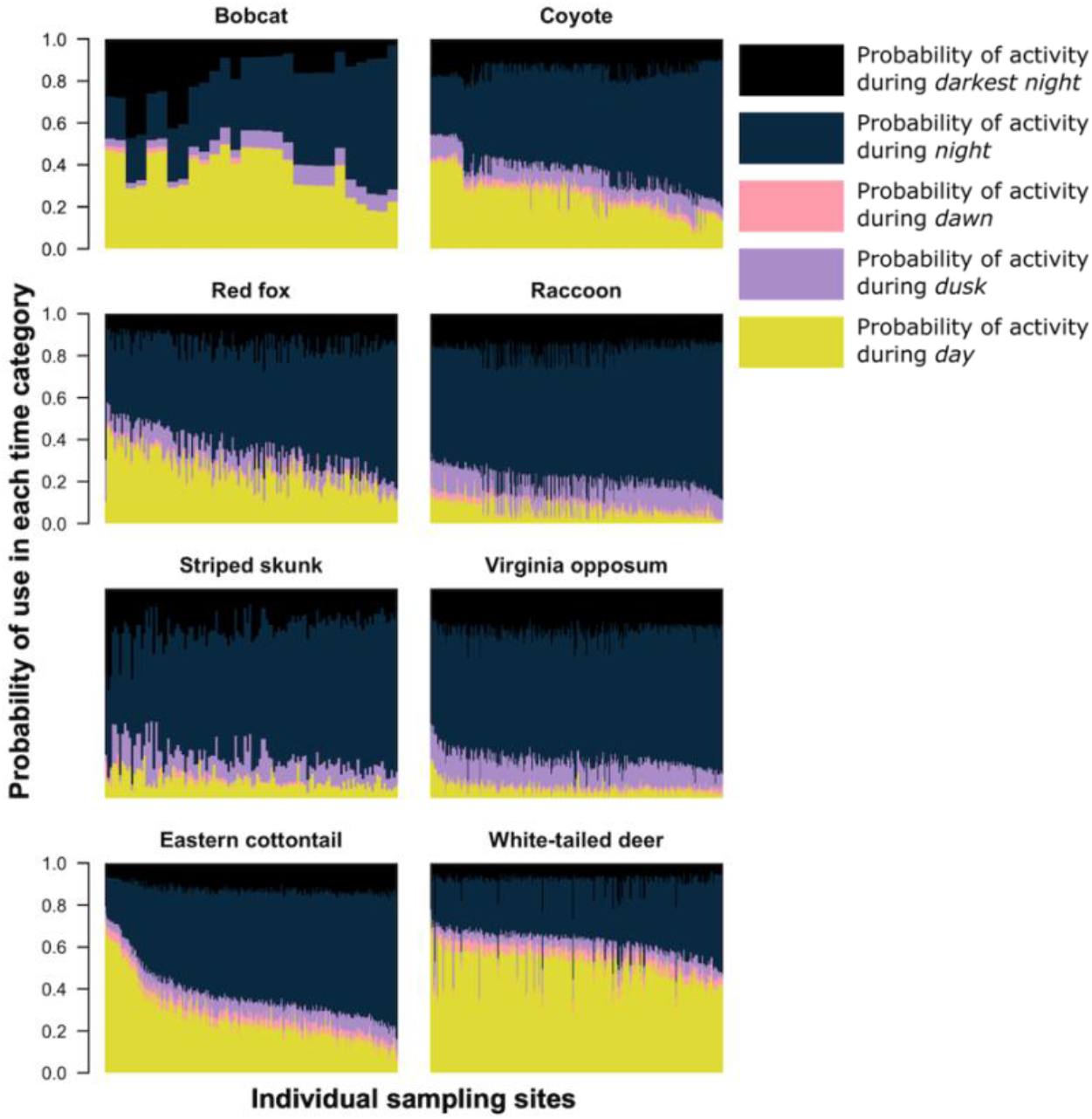
The predicted probability of activity in each time category at each sampling site (x-axis) the species was detected. Each column on the x-axis is a stacked bar plot representing the probability of activity in each time category at each sampling site. For each bar plot, all categories sum to one. Sampling sites along the x-axis are ordered from the lowest probability of nocturnal activity to the highest.

### Selection for particular time categories

Of the three predator species that we analyzed (coyote, bobcat, and red fox), we found that anthropogenic and natural features were associated with variation in diel activity for only coyote and red fox (Fig. 3a,b,c). Coyote selected for both nocturnal and crepuscular hours in areas of greater human population densities (Fig. 3b), and red fox avoided nocturnal hours in areas with more available greenspace (Fig. 3c). Seasonality also had an effect on both coyote and fox diel activity. Coyote selected for dawn hours (Fig. 3b) and red foxes selected for dusk hours during periods of higher daily average temperatures (Fig. 3c). We found no evidence that bobcats varied their diel activity across our environmental variables (Fig. 3a).

**Figure 3.**
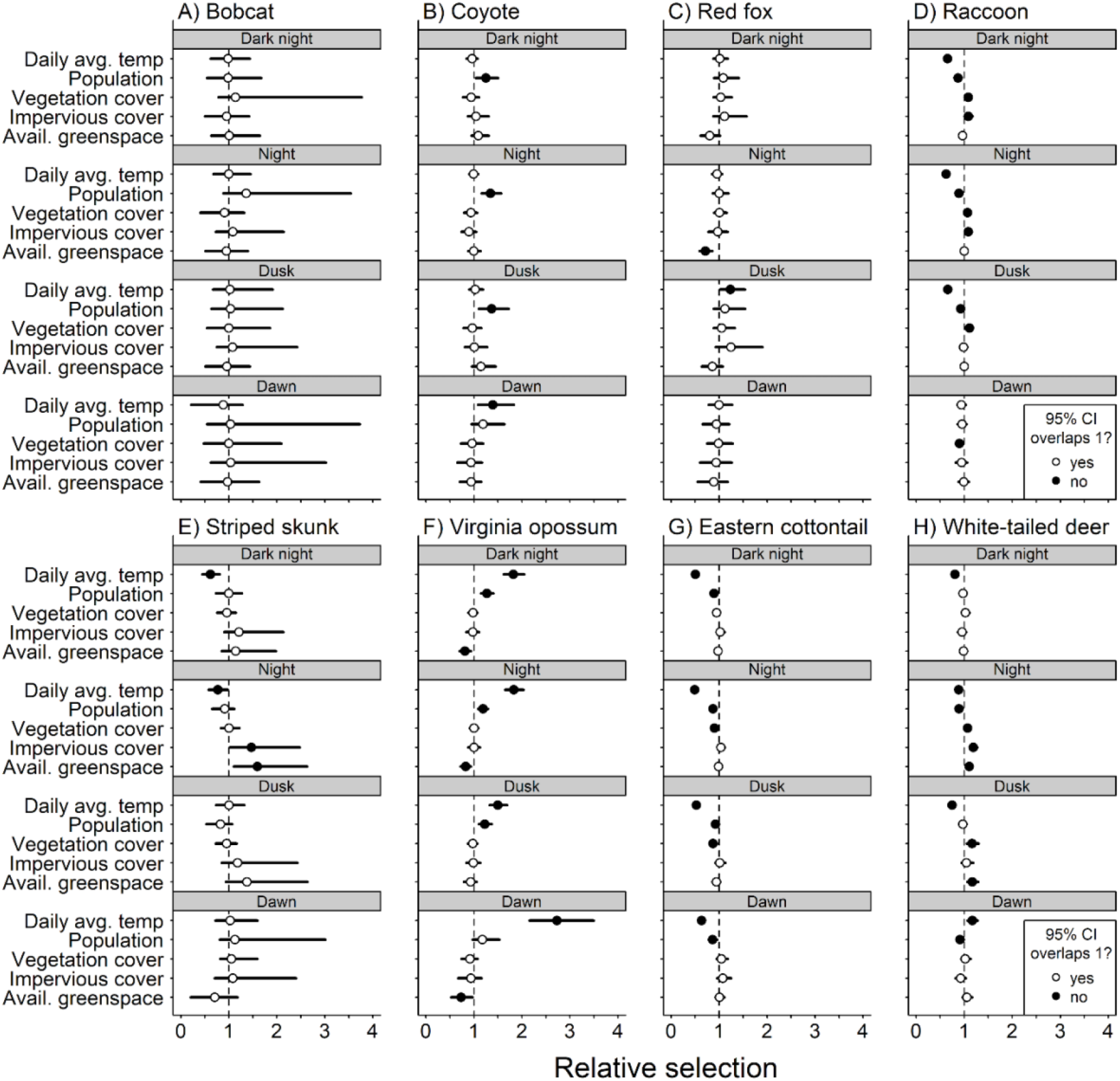
Mean (circle) and 95% credible intervals of estimated coefficients from natural and anthropogenic features on temporal selection of dark night, night, dusk, and dawn relative to day.

We found diel activity for all omnivore and herbivore species was affected by anthropogenic features. Raccoon, eastern cottontail, and white-tailed deer avoided nighttime hours in areas of greater human population density (Fig 3d,g,h), whereas Virginia opossum selected for nighttime and dusk hours in areas with greater human densities (Fig. 3f). Raccoon, striped skunk, and white-tailed deer all selected for nighttime hours in areas with greater impervious land cover (Fig. 3d,e,h).

Natural features were also associated with variation in diel activity for omnivore and herbivore species. As vegetation cover increased, eastern cottontails were more likely to select daytime hours (Fig. 3g), whereas raccoons and white-tailed deer were more likely to select for nighttime hours and dusk (Fig. 3d,h). As available greenspace increased, striped skunk were more likely to select nighttime hours (Fig. 3e), whereas Virginia opossum were less likely to select nighttime and dawn hours (Fig. 3f). White-tailed deer were also more likely to select nighttime and dusk hours as available greenspace increased (Fig. 3h).

We found seasonality effects on all omnivore and herbivore species. Virginia opossum were more likely to avoid daytime hours as temperatures increased (Fig. 3f). Daily average temperature had a positive relationship with diurnal selection for raccoons, striped skunk, and white-tailed deer (Fig. 3,d,e,h). Eastern cottontails, however, were more likely to select crepuscular hours and nighttime hours as daily average temperatures increased (Fig. 3g).

### Probability of nocturnal activity

Across all species, the probability of dawn and dusk activity was low (Figure 2 and 3). Therefore, we report the probability of nocturnal activity for each species by combining the probability of activity during night and darkest night. Coyote had a lower probability of being nocturnal in areas with lower human densities, but that probability increased significantly as human population increased (Fig. 4). With a one standard deviation (hereafter sd) increase from the mean human population density (from 1,512 – 3,095 people/km^2^), coyotes are 19% more likely to use nighttime hours and 38% more likely with a two sd increase from 1,512 to 4,678 people/km^2^ (Table 2). Red fox was the only species that had a significant change in the probability of nocturnal use across the available greenspace gradient (Fig. 4). Red fox were 23% less likely to use nighttime hours with a one sd increase in available greenspace from 0.41 to 0.57, and 41% less likely with a sd increase from 0.41 to 0.73 (Table 2). Note that predictor values vary because they were collected at species-specific scales and not all species were detected at the same sites.

**Figure 4.**
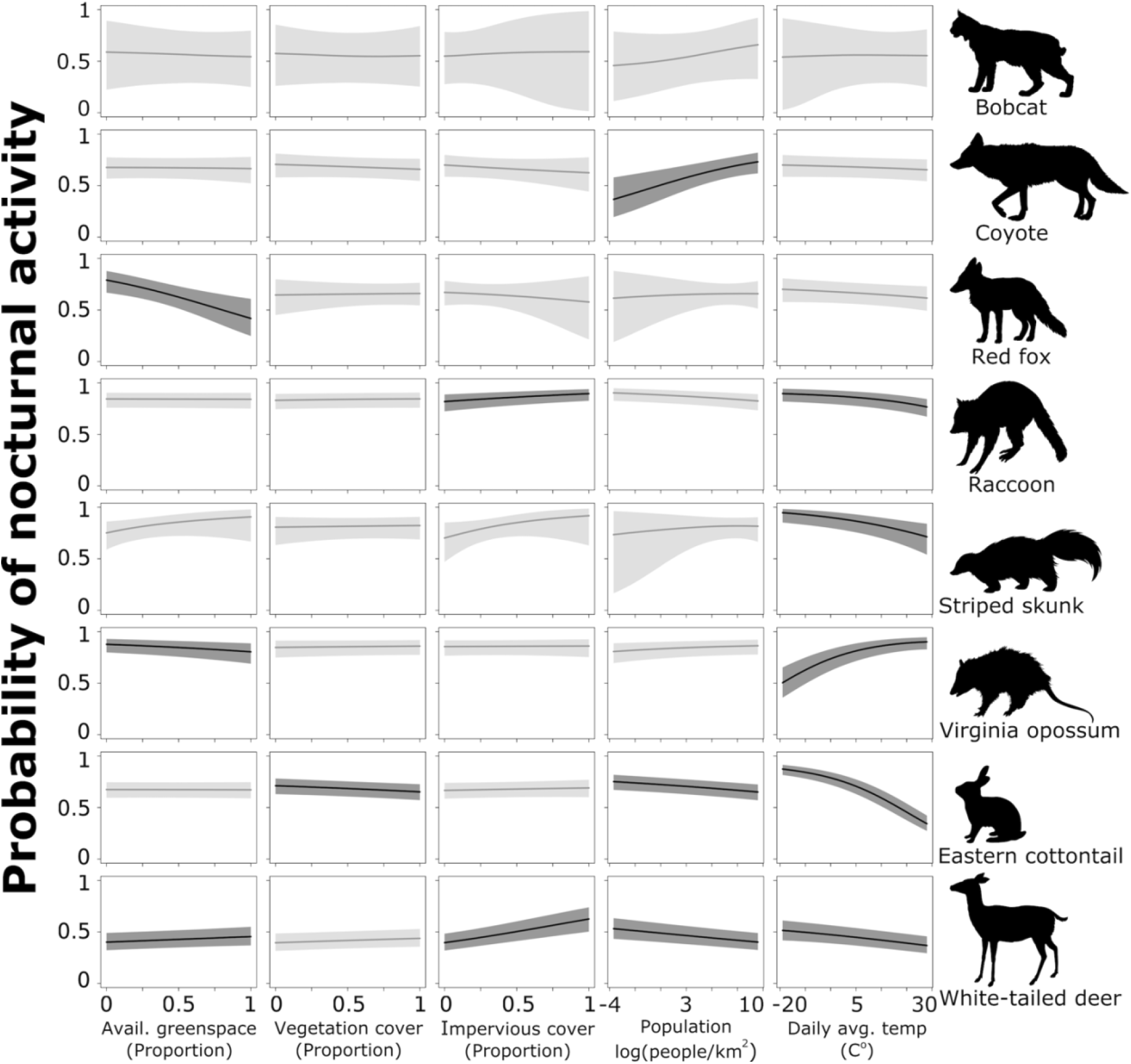
Probability of nocturnal activity (night and dark-night combined) across each of our natural and anthropogenic characteristics of the urban environment. Solid line indicates the median predicted line and shaded areas are 95% credible interval. Darker shading represent the relationships whose odds ratios did not overlap 1.

**Table 2.** Odds ratios for each predictor variable and a one and two standard deviation increase across their values. Bolded text indicates scenarios where the 95% credible intervals do not overlap 1.

|  | Available greenspace |  | Impervious cover |  | Vegetation Cover |  | Human pop. density |  | Daily avg. temp. |  |
| --- | --- | --- | --- | --- | --- | --- | --- | --- | --- | --- |
|  | 1-unit increase | 2-unit increase | 1-unit increase | 2-unit increase | 1-unit increase | 2-unit increase | 1-unit increase | 2-unit increase | 1-unit increase | 2-unit increase |
| Bobcat | 0.97 (0.62-1.38) | 0.95 (0.40-1.95) | 1.03 (0.70-1.76) | 1.06 (0.46-3.16) | 0.98 (0.55-1.56) | 0.99 (0.36-3.00) | 1.22 (0.85-2.81) | 1.51 (0.72-8.28) | 0.99 (0.71-1.37) | 0.99 (0.49-1.89) |
| Coyote | 0.99 (0.88-1.11) | 0.97 (0.76-1.23) | 0.94 (0.80-1.08) | 0.88 (0.64-1.18) | 0.95 (0.82-1.06) | 0.90 (0.68-1.13) | **1.19 (1.04-1.36)** | **1.38 (1.05-1.81)** | 0.95 (0.87-1.03) | 0.89 (0.75-1.05) |
| Red fox | **0.77 (0.65-0.90)** | **0.59 (0.41-0.81)** | 0.95 (0.77-1.15) | 0.90 (0.58-1.33) | 1.01 (0.90-1.13) | 1.01 (0.80-1.28) | 1.00 (0.87-1.17) | 1.00 (0.74-1.36) | 0.92 (0.83-1.02) | 0.85 (0.68-1.03) |
| Raccoon | 1.00 (0.96-1.03) | 0.99 (0.92-1.07) | **1.10 (1.05-1.16)** | **1.21 (1.1-1.34)** | 1.01 (0.97-1.05) | 1.02 (0.94-1.10) | **0.94 (0.90-0.97)** | **0.88 (0.81-0.95)** | **0.82 (0.77-0.87)** | **0.65 (0.57-0.73)** |
| Striped skunk | 1.26 (0.93-1.76) | 1.55 (0.82-3.00) | 1.26 (0.92-1.83) | 1.56 (0.80-3.31) | 1.02 (0.86-1.21) | 1.03 (0.74-1.46) | 1.01 (0.79-1.22) | 0.99 (0.57-1.46) | **0.73 (0.58-0.90)** | **0.54 (0.34-0.81)** |
| Virginia opossum | **0.88 (0.81-0.96)** | **0.77 (0.65-0.92)** | 1.01 (0.91-1.12) | 1.02 (0.83-1.25) | 1.02 (0.95-1.09) | 1.04 (0.91-1.18) | 1.04 (0.97-1.12) | 1.08 (0.93-1.24) | **1.27 (1.15-1.38)** | **1.49 (1.16-1.77)** |
| Eastern cottontail | 1.00 (0.95-1.04) | 1.00 (0.91-1.08) | 1.02 (0.97-1.09) | 1.05 (0.94-1.18) | **0.93 (0.88-0.98)** | **0.86 (0.78-0.95)** | **0.91 (0.87-0.95)** | **0.82 (0.75-0.89)** | **0.57 (0.54-0.61)** | **0.31 (0.28-0.35)** |
| White-tailed deer | **1.05 (1.00-1.10)** | **1.11 (1.00-1.22)** | **1.14 (1.07-1.21)** | **1.3 (1.15-1.46)** | 1.04 (0.99-1.09) | 1.08 (0.98-1.18) | **0.92 (0.88-0.96)** | **0.85 (0.78-0.92)** | **0.88 (0.84-0.92)** | **0.77 (0.71-0.84)** |

White-tailed deer, eastern cottontail, and raccoon had a greater probability of being active at night where human densities were low; this probability decreased as human population increased (Fig. 4). White-tailed deer were 8% less likely to use nighttime hours with a one sd increase in population density from 1,515 to 3,003 people/km^2^, eastern cottontail were 9% less likely (from 2,226 to 4,633 people/km^2^), and raccoon were 16% less likely (from 1,763 to 3,789 people/km^2^; Table 2). With a two sd increase in impervious cover (1,515 to 4,491 people/km^2^ for white-tailed deer, 2,226 to 7,040 for eastern cottontail, and 1,763 to 5,815 for raccoon), white-tailed deer were 16% less likely to be nocturnal, eastern cottontail 18% less likely, and raccoon 12% less likely to be nocturnal (Table 2). Conversely, white-tailed deer and raccoon showed a positive relationship with increased impervious cover and nocturnality (Fig. 4). White-tailed deer were 13% more likely to be active at night with a one sd increase in impervious cover from 0.16 to 0.31 and 29% more likely with a two sd increase from 0.16 to 0.45 (Table 2). Raccoons were 10% more likely to be active at night with a one sd increase in impervious cover and 21% more likely with an a two sd increase (Table 2).

Vegetation cover had a negative effect on the probability of nocturnal behavior of eastern cottontail (Fig. 4). Cottontail were 7% less likely to be nocturnal when the proportion of vegetation cover increased one sd above the mean from 0.67 to 0.92, and 14% less likely to be nocturnal when vegetation cover increased two sd above the mean from 0.67 to 1 (Table 2). We also found that white-tailed deer were 5% more likely to use nighttime hours when the proportion of available greenspace increased one sd above the mean from 0.52 to 0.75)and 11% more likely with an increase of two sd from the mean from 0.52 to 0.98 (Table 2). However, Virginia opossum were 12% less likely to be nocturnal with a one sd increase in available greenspace from 0.34 to 0.57 and 23% less likely with an increase of two sd from 0.34 to 0.78 (Table 2).

Finally, we found an influence of daily average temperature (season) on eastern cottontail, raccoon, striped skunk, white-tailed deer, and Virginia opossum (Fig. 4). Eastern cottontail were 43% less likely to use nighttime hours with a one sd increase in daily average temperature from 8.17 C to 18.76 C, and 69% less likely with a two sd increase from 8.17 C to 29.36 C (Table 2). With a one sd increase in temperature from 12.00 C to 21.49 C, raccoon were 18% less likely to use nighttime hours, and 35% less likely with a two sd increase from 12.00 C to 30.99 C (Table 2). Striped skunk were 27% less likely to exhibit nocturnal behavior with a one sd increase in daily average temperature from 15.3 C to 24.62 C, and 46% less with a two sd increase from 15.3 C to 33.94 C (Table 2). White-tailed deer were 12% less likely with a one sd increase from 12.01C to 22.65 C and 23% less likely with a two sd increase from 12.01 C to 33.30 C (Table 2). Virginia opossum, however, were 27% more likely to use nighttime hours with a one sd increase in daily average temperature from 13.85 C to 22.28 C, and 49% more likely with a two sd increase from 13.85 C to 30.71 C (Table 2). Again, temperature ranges vary because not all species were detected at the same sites and same times.

## Discussion

Ecological processes act across both space and time. We have, however, only just begun to study how animals use diel-time as an ecological resource to avoid risk and adapt to environmental change. We quantified the diel behavior of eight mammal species across urban gradients in ten U.S. cities. Our findings indicated that mammals can modulate their use of time within the 24-hour diel period as a resource to persist in urban ecosystems. We found that nocturnal activity had the greatest response to urbanization and seasonality, and that changes in nocturnality in response to urbanization were species-specific and varied among cities. Our results also illustrated the complex trade-offs that urban wildlife species must make to contend with both interspecific interactions (i.e., predation and competition) and human activity. These findings offer insight into how mammals might use time as a resource to adapt and persist in urban ecosystems.

We found that coyote had a greater probability of nocturnal behavior in areas with greater human densities. These findings are in agreement with past studies from single cities that documented increases in coyote nocturnal behavior in areas of higher human activity (Gallo et al., 2019; M. I. Grinder and Krausman, 2001; Riley et al., 2003; Tigas et al., 2002). Notably, vehicular collisions are a major mortality factor for coyotes (M. Grinder and Krausman, 2001) and coyotes have been typically persecuted by humans when they come in close contact (Dunlap, 1988; Young et al., 2019). Thus, a shift to nocturnal activity when traffic volumes are usually lower and humans are less active outdoors may be particularly important to survival in urban landscapes (Murray and St. Clair, 2015).

Red fox became less nocturnal as the proportion of local greenspace (i.e., available habitat) increased, a finding which may be explained by competition with coyote. Coyote and red fox exhibit a clear dominance hierarchy, whereby the dominant coyote negatively affects the subordinate red fox via competition and predation (Gosselink et al., 2003). Research has shown that urban coyotes occupy larger areas of greenspace (Gehrt et al., 2009). When more greenspace is available around a site, and presumably a higher probability of coyote presence, red foxes may become more diurnal to temporally avoid coyotes and reduce the risk of an interaction. Yet, when greenspace is limited, and presumably there is a lower probability of coyote presence, red foxes could be more active during nighttime hours with less risk of an interaction.

In most cases, the human aspects of urban environments captured by our predictor variables had opposite effects on omnivores and herbivores. While human population densities increased nocturnal activity for coyote, it decreased nocturnal activity for white-tailed deer, eastern cottontail, and raccoon. Prey species are known to spatially distribute themselves near human activity to act as a shield from predators (Berger, 2007; Shannon et al., 2014). In these cases, prey species may also utilize time as a human-mediated shield, exhibiting more activity at times of high human activity (daytime) in areas of high human densities. These results may seem counterintuitive given that increasing impervious cover increased the probability of nocturnal behavior exhibited by deer and raccoon (Fig. 4) and selection for nighttime hours by striped skunk (Fig. 3). However, a majority of impervious surfaces in the U.S. are roads and parking lots– places of high vehicular traffic (Frazer, 2005). Similar to coyote, vehicular collisions are a major source of mortality for these species (Glista et al., 2009). Therefore, a shift to nocturnal activity in areas with high impervious cover may be particularly important to their survival in cities and a sign of fine-scale modulation of temporal selection based on local environments.

Similarly, raccoon and white-tailed deer selected more for nighttime hours (Fig. 3) in locations with high levels of vegetation cover. More vegetation equates to more protective cover. Therefore, we suggest that raccoons and white-tailed deer can use the same temporal habitat as their predators (i.e., coyote) – but with less risk – when there is more physical cover. On the other hand, eastern cottontail were more diurnal with increased vegetation (Fig. 3 and 4), suggesting that more vegetation cover provides shelter from other perceived threats (i.e., humans; 22) and may allow eastern cottontail to select periods of high human activity (i.e. day). Interactions between various urban characteristics, which we did not examine in this study, should be further explored to fully understand how these characteristics jointly influence the temporal patterns of urban wildlife species.

Our results highlight the complexity of trade-offs for urban wildlife. In most cases, we found diverging activity patterns between coyote (a common urban apex predator) and subordinate or prey species in response to physical characteristics of urban environments. To persist in urban environments, it appears that urban species may have to modulate behaviors to contend with both anthropogenic risks and risk from predation or competition. Our results add to a growing body of literature that indicate species interactions in human-dominated landscapes may be better understood by explicitly considering the role humans play in those interactions (Berger, 2007; Blecha et al., 2018; Gallo et al., 2019; Magle et al., 2014).

We also found evidence that local climate, specifically temperature, regulated the diel behavior of many species. For example, white-tailed deer, eastern cottontail, and striped skunk became more diurnal as temperatures increased, presumably foraging more during the day in warmer seasons when more vegetation biomass is available. Virginia opossum showed a decrease in nocturnal behavior at lower temperatures. Given their poor thermoregulation abilities, poorly insulated fur, and cold-sensitive hands, ears, and tails (Kanda, 2005), it seems likely that Virginia opossum are morphologically constrained and thus unable to alter their diel activity patterns at colder temperatures. These results call attention to the importance of considering the impacts of morphology, physiology, and life history on a species’ capacity to adapt to environmental change. Given the interacting effects of climate change and urbanization (Stone, 2012), future research should explore how life history traits mediate temporal distributions of species activity – particularly as cities are rapidly warming (Oleson et al., 2015).

We did not find changes in diel activity for some species in response to our predictor variables. These results could be due to a lack of data on a particular species (i.e., bobcat) or because we did not sample across a large enough urban-rural gradient. Remote regions were not sampled in our study design, and some species may change their behavior at a lower level of urban intensity that we did not sample. Combining datasets from more rural and remote areas (e.g., Snapshot Serengeti (Swanson et al., 2015), Snapshot USA (Cove et al., 2021)) could allow us to identify the level of human development that elicits changes in diel activity for potentially sensitive species. Finally, our analysis was limited to the physical characteristics of cities. Additional characteristics like chronic noise, light pollution, resource supplementation, and species interactions influence animal behaviors and should be explored in future research.

Resource selection functions have been a popular and valuable tool to measure the relationship between available resources and animal populations, and have been used intensely in wildlife management and conservation (Strickland and McDonald, 2006). However, very little work has been done to quantify temporal habitat selection specifically (Cox et al., 2021; Gaston, 2019). Here, we built upon Farris et al. (Farris et al., 2015) and developed an analytical approach to quantify temporal resource selection across environmental gradients. While we have developed an analytical tool to measure temporal selection, a theoretical context for temporal habitat selection is needed and a further understanding of disproportional selection relative to the number of hours available is a promising avenue for future animal biology research.

Temporal partitioning may facilitate human-wildlife coexistence and effectively increase available habitats for species in cities. Temporal partitioning may also limit contact between people and animals, potentially reducing negative encounters like disease transmission and attacks on people (Gaynor et al., 2018). From a management perspective, ignoring diel behavior can result in biased estimates of species abundance and patterns of habitat use and lead to misinformed conservation measures (Gaston, 2019). Additionally, recognizing plasticity in species behavior can lead to better predictions of vulnerability to anthropogenic disturbances (Gaynor et al., 2018). Therefore, we recommend that diel activity and temporal partitioning be considered in conservation and management approaches.

We have shown that mammals have significant variation in the use and selection of time throughout the diel period. Additionally, our approach allowed us – for the first time – to quantify changes in diel activity across gradients of environmental change and across multiple urban areas, revealing that changes in diel patterns are influenced by natural and human landscape characteristics. Our results highlight the need to understand how a larger proportion of the animal community responds to urbanization, and provide evidence of behavioral plasticity that allows some species to adapt to and persist in human-dominated systems.

## Materials and Methods

### Study Design

The number of sampling sites per city ranged from 24-113 (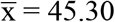, sd = 28.65). In each city, sampling sites were placed along a gradient of urbanization (high to low population density and impervious cover). At each sampling site (*n =* 453) we placed one Bushnell motion-triggered infrared Trophy Cam (Bushnell Corp., Overland Park, KS, USA). Sampling sites were located in greenspaces, such as city parks, cemeteries, natural areas, utility easements, and golf courses. To increase the detection probability of each species we placed one synthetic fatty acid scent lure in the camera line of sight, and lures were replaced on two-week intervals if missing to remain consistent throughout the study. However, Fidino et al. (2020) later found that this type of lure has little to no effect on the detectability of most urban mammals. We used observation data collected between January 2017 and December 2018. However, not all cities were sampled continuously throughout the study period (Table S1).

### Data processing

For each species, we defined a single detection event as all photos taken within a 15-minute period at each camera station (Farris et al., 2015; Ridout and Linkie, 2009). We categorized each detection event as either ‘dawn’, ‘dusk’, ‘day’, ‘night’, and ‘darkest night’ using the *suncalc* package (Thieurmel and Elmarhraoui, 2019) in R ver 4.2.0 (R Core Team, 2019). The suncalc package defines and calculates ‘dawn’ as starting when morning astronomical twilight begins and ending when the bottom edge of the sun touches the horizon. ‘Dusk’ was defined as the beginning of evening twilight to the point when it became dark enough for astronomical observations. ‘Day’ was defined as the period between dawn and dusk. We considered the nighttime as two distinct time periods (night and darkest night), because some species may be nocturnal but use the darkest hours of the night to reduce the risk of human interactions (Gehrt et al., 2009). We defined ‘night’ as the periods between the end of dusk and one hour before the darkest moment of the night (when the sun is at the lowest point), and from one hour after the darkest moment to dawn. The ‘darkest hours’ of the night were categorized as one hour before and after the darkest moment in the night. We accounted for the date, geographical location, and daylight savings time of each detection events. Therefore, the amount of time available in each category could vary geographically and seasonally.

### Predictor Variables

To assess how characteristics of urban environments influenced diel activity of urban wildlife mammals, we calculated site-level predictor variables within a fixed-radius buffer around each sampling site. Fixed-radius buffers varied in size among species and were based on the typical home range of each species: 500 m fixed-radius buffer for eastern cottontail (Hunt et al., 2014), Virginia opossum (Fidino et al., 2016; Wright et al., 2012), and white-tailed deer (Etter et al., 2002); 1 km fixed-radius buffer for striped skunk (Weissinger et al., 2009) and raccoon (Rosatte, 2000), and 1.5 km fixed radius buffer for coyote (Gehrt et al., 2009; Riley et al., 2003), red fox (Mueller et al., 2018), and bobcat (Riley et al., 2003). In our analysis, we included variables that described two contrasting characteristics of urban ecosystems, the natural and the human-built environment (Table S4). We also included average temperature to account for possible seasonal changes in diel activity.

#### Urban features

To characterize urbanization around each sampling site, we calculated human population density (individuals/km^2^) and mean impervious cover (%). Population density was extracted from Block Level Housing Density data (Radeloff et al., 2018) created from 2010 U.S. Census data (U.S. Census Bureau, 2010). Mean impervious cover was calculated from the 2011 National Land Cover Database (NLCD) 30-m resolution Percent Developed Imperviousness data (Homer et al., 2015).

#### Natural features

To characterize natural features, we calculated the proportion of vegetation cover and the proportion of available greenspace (i.e., potential habitat) around each site. To calculate the proportion of vegetation cover around each sampling site, we first calculated the Normalized Difference Vegetation Index (NDVI) using U.S. Geological Survey 30-m resolution LandSat 8 data that 1) covered the entire study area of each city, 2) was taken during a summer month that coincided with the respective city’s sampling period, and 3) contained less than 15% cloud cover. LandSat 8 imagery was downloaded with R using the *getSpatialData* package (Schwalb-Willmann, 2019). We then calculated vegetation cover as the proportion of cells within each fixed-radius buffer that had an NDVI value representing substantial vegetation cover (> 0.2; https://climatedataguide.ucar.edu/climate-data/ndvi-normalized-difference-vegetation-index-noaa-avhrr). To calculate available greenspace, we extracted the proportion of 2011 NLCD Land Cove 30-m resolution raster cells within each fixed-radius buffer that were classified as forest, shrubland, herbaceous, wetland, and developed open space (which included urban green spaces).

#### Seasonality

Because weather that defines each calendar season varies across our sampled longitudinal gradient, we used daily average temperature (i.e., mean temperature on the day of a given detection event) as a continuous covariate to describe seasonality. For each day and location of a detection event, we recorded the daily average temperature from the National Climatic Data Center using the R package *rnoaa* (Chamberlain, 2020). We used data from the nearest weather station to each city that recorded daily weather during our study period (Table S5).

#### Quantifying the influence of urban characteristics on diel patterns

By splitting diel time into *k* in 1,…,K categories, we estimated the probability a detection event occurs in each category for each species. To do so, we let *y_i_* be the time category of the *i*^th^ in 1,…,*I* detection events, and assume it is a Categorical random variable, where ***ϕ*** is a probability vector of the K categories ***ϕ*** = [*ϕ*_1_*ϕ*_2_*ϕ*_3_*ϕ*_4_*ϕ*_5_], *ϕ*_1_ = 1 − *ϕ*_2_ − *ϕ*_3_ − *ϕ*_4_ − *ϕ*_5_, and **1**⋅ ***ϕ*** = 1 such that:

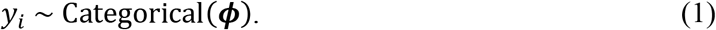

To understand mechanistic changes in species-specific diel activity patterns and assess the influence that each predictor variable had on the temporal activity of each species, we let ***ϕ***_***i***_ be a function of covariates with the softmax function,

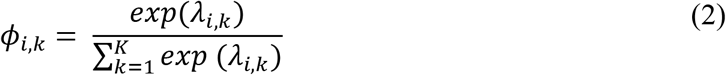

where *λ*_*i*,*k*_ is the log-linear predictor for detection event *i* and category *k*. We set our reference category as ‘day’ (i.e., *k* = 1). In our model the log-linear predictor of each outcome is then

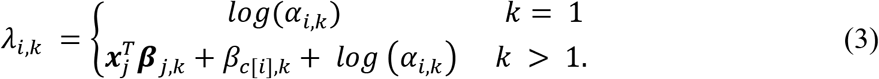

In Eq. 3, ***β**_j,k_* coefficients correspond to the effect of greenspace availability, impervious cover, vegetation cover, human population density, and daily average temperature for k > 1. As detection events within each city may not be wholly independent, we included a random intercept for city, *β_c[i],k_*, where **c** is a vector of length *I* that denotes which city detection event *i* occurred (Gelman and Hill, 2006). Finally, to account for the different amount of time available to animals among the *K* categories, we also included a log offset term, log(α_*k,i*_), where α_*k,i*_ is the number of hours available in category *k* at the time of detection event *i*. This form of multinomial regression is equivalent to a logistic regression model with a spatial categorical covariate with K levels, where the offset accounts for varying availability. As such, our model approximates the weighted distribution used in resource selection functions assuming an exponential link (Hooten et al., 2017). Exponentiated coefficient estimates greater than one indicates ‘selection’ and less than one indicates ‘avoidance’, relative to the day reference category.

Because we considered ‘day’ (k = 1) as our reference outcome, we set β_*c[i]*,1_ = 0 and β_*j*,1_ = 0 (Eq. 3). The remaining **β**_*j,k*_ parameters were given Laplace(0,π) priors as a form of categorical LASSO regularization (Tutz et al., 2015). We took a fully Bayesian approach to variable selection by estimating the hyperparameter π (van Erp et al., 2019), which was given a uniform(0.001,10) prior distribution. β_c[i],k_ was given a Normal(μ_*k*_,τ_*k*_) prior for each city where μ_*k*_ ~ Normal(0,10) and τ_*k*_ ~ Gamma(1,1).

Models were fit using an Markov Chain Monte Carlo (MCMC) algorithm implemented in JAGS ver 4.2.0 (Plummer, 2003) using the *runjags* package (Denwood, 2016) in R. Fourteen parallel chains were each run from random starting values. The first 20,000 iterations from each chain were discarded and every 7^th^ iteration was kept to reduce autocorrelation among the samples. A total of 75,000 iterations were obtained for each model. Model convergence was assessed by checking that the Gelman-Rubin diagnostic statistic for each parameter was <1.1 (Gelman and Rubin, 1992) and by visually inspecting the trace plots of MCMC sample

## Acknowledgements

The authors would like to thank all the field technicians, students, and assistants associated with the Urban Wildlife Information Network for data collection and photo processing. We would also like to thank the operations, facilities, and administrative staff at our respective institutions as their work behind the scenes is vital to our research. Funding was provided by the Abra Prentice-Wilkin Foundation and the EJK Foundation. We would also like to thank N. Clemente, J. Kimlinger, and Pariveda Solutions for their help with an application to store and tag our camera trap images.

## Competing Interest Statement

No Competing Interests

## Notes

### Competing Interest Statement

The authors have declared no competing interest.

## References

Berger J. 2007. Fear, human shields and the redistribution of prey and predators in protected areas. Biol Lett 3:620–623. doi:10.1098/rsbl.2007.0415

Blecha KA, Boone RB, Alldredge MW. 2018. Hunger mediates apex predator’s risk avoidance response in wildland–urban interface. J Anim Ecol 87:609–622. doi:10.1111/1365-2656.12801

Chamberlain S. 2020. rnoaa: “NOAA” Weather Data from R. R package version 0.9.6.

Clinchy M, Zanette LY, Roberts D, Suraci JP, Buesching CD, Newman C, Macdonald DW. 2016. Fear of the human “super predator” far exceeds the fear of large carnivores in a model mesocarnivore. Behav Ecol 27:1826–1832. doi:10.1093/beheco/arw117

Cove MV, Kays R, Bontrager H, Bresnan C, Lasky M, Frerichs T, Klann R, Lee Jr. TE, Crockett SC, Crupi AP, Weiss KCB, Rowe H, Sprague T, Schipper J, Tellez C, Lepczyk CA, Fantle-Lepczyk JE, LaPoint S, Williamson J, Fisher-Reid MC, King SM, Bebko AJ, Chrysafis P, Jensen AJ, Jachowski DS, Sands J, MacCombie KA, Herrera DJ, van der Merwe M, Knowles TW, Horan III RV, Rentz MS, Brandt LSE, Nagy C, Barton BT, Thompson WC, Maher SP, Darracq AK, Hess G, Parsons AW, Wells B, Roemer GW, Hernandez CJ, Gompper ME, Webb SL, Vanek JP, Lafferty DJR, Bergquist AM, Hubbard T, Forrester T, Clark D, Cincotta C, Favreau J, Facka AN, Halbur M, Hammerich S, Gray M, Rega-Brodsky CC, Durbin C, Flaherty EA, Brooke JM, Coster SS, Lathrop RG, Russell K, Bogan DA, Cliché R, Shamon H, Hawkins MTR, Marks SB, Lonsinger RC, O’Mara MT, Compton JA, Fowler M, Barthelmess EL, Andy KE, Belant JL, Beyer Jr. DE, Kautz TM, Scognamillo DG, Schalk CM, Leslie MS, Nasrallah SL, Ellison CN, Ruthven C, Fritts S, Tleimat J, Gay M, Whittier CA, Neiswenter SA, Pelletier R, DeGregorio BA, Kuprewicz EK, Davis ML, Dykstra A, Mason DS, Baruzzi C, Lashley MA, Risch DR, Price MR, Allen ML, Whipple LS, Sperry JH, Hagen RH, Mortelliti A, Evans BE, Studds CE, Sirén APK, Kilborn J, Sutherland C, Warren P, Fuller T, Harris NC, Carter NH, Trout E, Zimova M, Giery ST, Iannarilli F, Higdon SD, Revord RS, Hansen CP, Millspaugh JJ, Zorn A, Benson JF, Wehr NH, Solberg JN, Gerber BD, Burr JC, Sevin J, Green AM, Şekercioğlu ÇH, Pendergast M, Barnick KA, Edelman AJ, Wasdin JR, Romero A, O’Neill BJ, Schmitz N, Alston JM, Kuhn KM, Lesmeister DB, Linnell MA, Appel CL, Rota C, Stenglein JL, Anhalt-Depies C, Nelson C, Long RA, Jo Jaspers K, Remine KR, Jordan MJ, Davis D, Hernández-Yáñez H, Zhao JY, McShea WJ. 2021. SNAPSHOT USA 2019: a coordinated national camera trap survey of the United States. Ecology 102:e03353. doi:10.1002/ecy.3353

Cox DTC, Gardner AS, Gaston KJ. 2021. Diel niche variation in mammals associated with expanded trait space. Nat Commun 12:1753. doi:10.1038/s41467-021-22023-4

Denwood MJ. 2016. runjags: An R Package Providing Interface Utilities, Model Templates, Parallel Computing Methods and Additional Distributions for MCMC Models in JAGS. J Stat Softw 71:25.

Dunlap T. 1988. Saving America’s Wildlife: Ecology and the American Mind, 1850-1990. New Jersey, USA: Princeton University Press.

Etter DR, Hollis KM, Van Deelen TR, Ludwig DR, Chelsvig JE, Anchor CL, Warner RE. 2002. Survival and Movements of White-Tailed Deer in Suburban Chicago, Illinois. J Wildl Manag 66:500–510. doi:10.2307/3803183

Farris ZJ, Gerber BD, Karpanty S, Murphy A, Andrianjakarivelo V, Ratelolahy F, Kelly MJ. 2015. When carnivores roam: temporal patterns and overlap among Madagascar’s native and exotic carnivores. J Zool 296:45–57. doi:10.1111/jzo.12216

Fidino M, Barnas GR, Lehrer EW, Murray MH, Magle SB. 2020. Effect of Lure on Detecting Mammals with Camera Traps. Wildl Soc Bull 44:543–552. doi:10.1002/wsb.1122

Fidino M, Gallo T, Lehrer EW, Murray MH, Kay CAM, Sander HA, MacDougall B, Salsbury CM, Ryan TJ, Angstmann JL, Amy Belaire J, Dugelby B, Schell CJ, Stankowich T, Amaya M, Drake D, Hursh SH, Ahlers AA, Williamson J, Hartley LM, Zellmer AJ, Simon K, Magle SB. 2021. Landscape-scale differences among cities alter common species’ responses to urbanization. Ecol Appl 31:e02253. doi:10.1002/eap.2253

Fidino MA, Lehrer EW, Magle SB. 2016. Habitat Dynamics of the Virginia Opossum in a Highly Urban Landscape. Am Midl Nat 175:155–167. doi:10.1674/0003-0031-175.2.155

Frazer L. 2005. Paving paradise: the peril of impervious surfaces. Environ Health Perspect 113:A456–A462. doi:10.1289/ehp.113-a456

Gallo T, Fidino M, Lehrer EW, Magle S. 2019. Urbanization alters predator-avoidance behaviours. J Anim Ecol 88:793–803. doi:10.1111/1365-2656.12967

Gaston KJ. 2019. Nighttime Ecology: The “Nocturnal Problem” Revisited. Am Nat 193:481–502.

Gaynor KM, Hojnowski CE, Carter NH, Brashares JS. 2018. The influence of human disturbance on wildlife nocturnality. Science 360:1232–1235. doi:10.1126/science.aar7121

Gehrt SD, Anchor C, White LA. 2009. Home Range and Landscape Use of Coyotes in a Metropolitan Landscape: Conflict or Coexistence? J Mammal 90:1045–1057.

Gelman A, Hill J. 2006. Data Analysis Using Regression and Multilevel/Hierarchical Models, 1 edition. ed. Cambridge ; New York: Cambridge University Press.

Gelman A, Rubin DB. 1992. Inference from iterative simulation using multiple sequences. Stat Sci 7:457–472.

Glista DJ, DeVault TL, DeWoody JA. 2009. A review of mitigation measures for reducing wildlife mortality on roadways. Landsc Urban Plan 91:1–7. doi:10.1016/j.landurbplan.2008.11.001

Gosselink TE, Van Deelen TR, Warner RE, Joselyn MG. 2003. Temporal Habitat Partitioning and Spatial Use of Coyotes and Red Foxes in East-Central Illinois. J Wildl Manag 67:90–103. doi:10.2307/3803065

Grinder M, Krausman P. 2001. Morbidity-mortality factors and survival of an urban coyote population in Arizona. J Wildl Dis 37.

Grinder MI, Krausman PR. 2001. Home Range, Habitat Use, and Nocturnal Activity of Coyotes in an Urban Environment. J Wildl Manag 65:887–898. doi:10.2307/3803038

Homer CG, Dewitz J, Yang L, Jin S, Danielson P, Xian, Coulston J, Herold N, Wickham J, Megown J. 2015. Completion of the 2011 National Land Cover Database for the conterminous United States –representing a decade of land cover change information. Photogramm Eng Remote Sens 81:345–353.

Hooten MB, Johnson DS, McClintock BT, Morales JM. 2017. Animal movement: statistical models for telemetry data. CRC press.

Hunt VM, Magle SB, Vargas C, Brown AW, Lonsdorf EV, Sacerdote AB, Sorley EJ, Santymire RM. 2014. Survival, abundance, and capture rate of eastern cottontail rabbits in an urban park. Urban Ecosyst 17:547–560. doi:10.1007/s11252-013-0334-z

Kanda LL. 2005. Winter energetics of Virginia opossums Didelphis virginiana and implications for the species’ northern distributional limit. Ecography 28:731–744. doi:10.1111/j.2005.0906-7590.04173.x

Kitchen AM, Gese EM, Schauster ER. 2000. Changes in coyote activity patterns due to reduced exposure to human persecution. Can J Zool 78:853–857. doi:10.1139/z00-003

Kronfeld-Schor N, Dayan T. 2003. Partitioning of Time as an Ecological Resource. Annu Rev Ecol Evol Syst 34:153–181. doi:10.1146/annurev.ecolsys.34.011802.132435

Kruschke JK. 2011. Doing Bayesian Data Analysis, 2nd ed. London, UK: Academic Press.

Lehrer EW, Gallo T, Fidino M, Kilgour RJ, Wolff PJ, Magle SB. 2021. Urban bat occupancy is highly influenced by noise and the location of water: Considerations for nature-based urban planning. Landsc Urban Plan 210:104063. doi:10.1016/j.landurbplan.2021.104063

Levy O, Dayan T, Porter WP, Kronfeld-Schor N. 2019. Time and ecological resilience: can diurnal animals compensate for climate change by shifting to nocturnal activity? Ecol Monogr 89:e01334. doi:10.1002/ecm.1334

Magle SB, Fidino M, Lehrer EW, Gallo T, Mulligan MP, Ríos MJ, Ahlers AA, Angstmann J, Belaire A, Dugelby B, Gramza A, Hartley L, MacDougall B, Ryan T, Salsbury C, Sander H, Schell C, Simon K, Onge SS, Drake D. 2019. Advancing urban wildlife research through a multi-city collaboration. Front Ecol Environ 17:232–239. doi:10.1002/fee.2030

Magle SB, Simoni LS, Lehrer EW, Brown JS. 2014. Urban predator–prey association: coyote and deer distributions in the Chicago metropolitan area. Urban Ecosyst 875–891. doi:10.1007/s11252-014-0389-5

Maloney SK, Moss G, Cartmell T, Mitchell D. 2005. Alteration in diel activity patterns as a thermoregulatory strategy in black wildebeest (Connochaetes gnou). J Comp Physiol A 191:1055–1064. doi:10.1007/s00359-005-0030-4

McDonnell MJ, Pickett STA. 1990. Ecosystem Structure and Function along Urban-Rural Gradients: An Unexploited Opportunity for Ecology. Ecology 71:1232–1237. doi:10.2307/1938259

Morey PS, Gese EM, Gehrt S. 2007. Spatial and Temporal Variation in the Diet of Coyotes in the Chicago Metropolitan Area. Am Midl Nat 158:147–161. doi:10.1674/0003-0031(2007)158[147:SATVIT]2.0.CO;2

Mueller MA, Drake D, Allen ML. 2018. Coexistence of coyotes (Canis latrans) and red foxes (Vulpes vulpes) in an urban landscape. PLOS ONE 13:e0190971. doi:10.1371/journal.pone.0190971

Murray MH, St. Clair CC. 2017. Predictable features attract urban coyotes to residential yards. J Wildl Manag 81:593–600. doi:10.1002/jwmg.21223

Murray MH, St. Clair CC. 2015. Individual flexibility in nocturnal activity reduces risk of road mortality for an urban carnivore. Behav Ecol 26:1520–1527. doi:10.1093/beheco/arv102

Oleson KW, Monaghan A, Wilhelmi O, Barlage M, Brunsell N, Feddema J, Hu L, Steinhoff DF. 2015. Interactions between urbanization, heat stress, and climate change. Clim Change 129:525–541. doi:10.1007/s10584-013-0936-8

Pacione Michael. 2009. Urban geography : a global perspective, 3rd ed. ed. London ; Routledge.

Plummer M. 2003. A program for analysis of Bayesian graphical models using Gibbs samplingProceedings of the Third International Workshop on Distributed Statistical Computing. Vienna, Austria: R Foundations for Statistical Computing. pp. 125–133.

R Core Team. 2019. R: A language and environment for statistical computing. Vienna, Austria: R Foundation for Statistical Computing.

Radeloff VC, Helmers DP, Kramer HA, Mockrin MH, Alexandre PM, Bar-Massada A, Butsic V, Hawbaker TJ, Martinuzzi S, Syphard AD, Stewart SI. 2018. Rapid growth of the US wildland-urban interface raises wildfire risk. Proc Natl Acad Sci 115:3314–3319. doi:10.1073/pnas.1718850115

Ridout MS, Linkie M. 2009. Estimating overlap of daily activity patterns from camera trap data. J Agric Biol Environ Stat 14:322–337. doi:10.1198/jabes.2009.08038

Riley SPD, Sauvajot RM, Fuller TK, York EC, Kamradt DA, Bromley C, Wayne RK. 2003. Effects of Urbanization and Habitat Fragmentation on Bobcats and Coyotes in Southern California. Conserv Biol 17:566–576. doi:10.1046/j.1523-1739.2003.01458.x

Rosatte RC. 2000. Management of raccoons (Procyon lotor) in Ontario, Canada: Do human intervention and disease have significant impact on raccoon populations? Mammalia 64:369–390. doi:10.1515/mamm.2000.64.4.369

Schoener TW. 1974. Resource Partitioning in Ecological Communities. Science 185:27. doi:10.1126/science.185.4145.27

Schwalb-Willmann J. 2019. getSpatialData: R package version 0.0.4.

Shannon G, Cordes LS, Hardy AR, Angeloni LM, Crooks KR. 2014. Behavioral Responses Associated with a Human-Mediated Predator Shelter. PLOS ONE 9:e94630. doi:10.1371/journal.pone.0094630

Stone B. 2012. The City and the Coming Climate: Climate Change in the Places We Live. Cambridge University Press.

Strickland MD, McDonald LL. 2006. Introduction to the Special Section on Resource Selection. J Wildl Manag 70:321–323.

Swanson A, Kosmala M, Lintott C, Simpson R, Smith A, Packer C. 2015. Snapshot Serengeti, high-frequency annotated camera trap images of 40 mammalian species in an African savanna. Sci Data 2:150026. doi:10.1038/sdata.2015.26

Tambling CJ, Minnie L, Meyer J, Freeman EW, Santymire RM, Adendorff J, Kerley GIH. 2015. Temporal shifts in activity of prey following large predator reintroductions. Behav Ecol Sociobiol 69:1153–1161. doi:10.1007/s00265-015-1929-6

Thieurmel B, Elmarhraoui A. 2019. suncalc: Compute Sun Position, Sunlight Phases, Moon Position and Lunar Phase.

Tigas LA, Van Vuren DH, Sauvajot RM. 2002. Behavioral responses of bobcats and coyotes to habitat fragmentation and corridors in an urban environment. Biol Conserv 108:299–306. doi:10.1016/S0006-3207(02)00120-9

Tutz G, Pößnecker W, Uhlmann L. 2015. Variable selection in general multinomial logit models. Comput Stat Data Anal 82:207–222. doi:10.1016/j.csda.2014.09.009

U.S. Census Bureau. 2010. Population, Housing Units, Area, and Density: 2010 - County -- County Subdivision and Place more information 2010 Census Summary File 1. Washington, D.C. USA: U.S. Department of Commerce.

van der Vinne V, Tachinardi P, Riede SJ, Akkerman J, Scheepe J, Daan S, Hut RA. 2019. Maximising survival by shifting the daily timing of activity. Ecol Lett 22:2097–2102. doi:10.1111/ele.13404

van Erp S, Oberski DL, Mulder J. 2019. Shrinkage priors for Bayesian penalized regression. J Math Psychol 89:31–50. doi:10.1016/j.jmp.2018.12.004

Weissinger MD, Theimer TC, Bergman DL, Deliberto TJ. 2009. NIGHTLY AND SEASONAL MOVEMENTS, SEASONAL HOME RANGE, AND FOCAL LOCATION PHOTO-MONITORING OF URBAN STRIPED SKUNKS (MEPHITIS MEPHITIS): IMPLICATIONS FOR RABIES TRANSMISSION. J Wildl Dis 45:388–397. doi:10.7589/0090-3558-45.2.388

Wolkovich EM, Cook BI, McLauchlan KK, Davies TJ. 2014. Temporal ecology in the Anthropocene. Ecol Lett 17:1365–1379. doi:10.1111/ele.12353

Wright JD, Burt MS, Jackson VL. 2012. Influences of an Urban Environment on Home Range and Body Mass of Virginia Opossums (Didelphis virginiana). Northeast Nat 19:77–86.

Young JK, Hammill E, Breck SW. 2019. Interactions with humans shape coyote responses to hazing. Sci Rep 9:20046. doi:10.1038/s41598-019-56524-6

