## Supplemental Materials for "Mammals adjust diel activity across gradients of urbanization"

Table S1. Camera trap start and end date, the total number of sampling sites, and total number of trap nights for each city used to analyze diel patterns in urban mammals.

| City | Start Date | End Date | Sites | Trap Nights |
|---|---|---|---|---|
| Austin, Texas | 3/28/17 | 8/7/17 | 25 | 612 |
| Chicago, Illinois | 1/9/17 | 11/11/18 | 113 | 14515 |
| Denver, Colorado | 1/1/17 | 7/29/17 | 40 | 2695 |
| Fort Collins, Colorado | 1/4/17 | 5/23/18 | 31 | 3597 |
| Iowa City, Iowa | 6/12/17 | 11/1/18 | 39 | 6912 |
| Indianapolis, Indiana | 1/3/17 | 2/9/18 | 55 | 5634 |
| Long Beach, California | 5/31/18 | 11/8/18 | 24 | 565 |
| Manhattan, Kansas | 6/28/17 | 9/1/17 | 74 | 1554 |
| Madison, Wisconsin | 1/12/17 | 7/10/18 | 24 | 3548 |
| Wilmington, Delaware | 2/9/18 | 11/1/18 | 28 | 1862 |

Table S2. The proportion of sampling sites in each city that each species was detected on camera between January 2017 and December 2018

| | Austin, TX | Chicago, IL | Denver, CO | Ft. Collins, CO | Iowa City, IA | Indianapolis, IN | Long Beach, CA | Manhattan, KS | Madison, WI | Wilmington, DE |
|---|---|---|---|---|---|---|---|---|---|---|
| Bobcat | 0.04 | 0.00 | 0.00 | 0.13 | 0.18 | 0.00 | 0.33 | 0.11 | 0.00 | 0.00 |
| Coyote | 0.36 | 0.81 | 0.58 | 0.58 | 0.87 | 0.80 | 0.79 | 0.43 | 0.42 | 0.00 |
| Eastern cottontail | 0.12 | 0.80 | 0.48 | 0.81 | 0.85 | 0.87 | 0.42 | 0.32 | 0.96 | 0.50 |
| Raccoon | 0.52 | 0.86 | 0.62 | 1.00 | 1.00 | 0.93 | 0.21 | 0.80 | 0.75 | 1.00 |
| Red fox | 0.00 | 0.09 | 0.32 | 0.58 | 0.82 | 0.62 | 0.00 | 0.16 | 0.50 | 1.00 |
| Striped skunk | 0.08 | 0.52 | 0.17 | 0.45 | 0.18 | 0.04 | 0.17 | 0.14 | 0.29 | 0.36 |
| Virginia opossum | 0.28 | 0.85 | 0.00 | 0.00 | 1.00 | 0.87 | 0.21 | 0.66 | 0.92 | 0.82 |
| White-tailed deer | 0.44 | 0.50 | 0.22 | 0.74 | 1.00 | 0.71 | 0.21 | 0.57 | 0.29 | 0.89 |

Table S3. Range of probabilities of activity in each time category across all sampling sites.

| Species | Category | Minimum probability | Maximum probability |
|---|---|---|---|
| *Bobcat* | Day | 0.18 | 0.49 |
| | Dawn | 0.00 | 0.03 |
| | Dusk | 0.02 | 0.10 |
| | Night | 0.20 | 0.69 |
| | Darkest night | 0.03 | 0.47 |
| *Coyote* | Day | 0.07 | 0.44 |
| | Dawn | 0.01 | 0.06 |
| | Dusk | 0.04 | 0.20 |
| | Night | 0.27 | 0.70 |
| | Darkest night | 0.09 | 0.24 |
| *Eastern cottontail* | Day | 0.05 | 0.72 |
| | Dawn | 0.02 | 0.06 |
| | Dusk | 0.02 | 0.11 |
| | Night | 0.15 | 0.69 |
| | Darkest night | 0.05 | 0.18 |
| *Raccoon* | Day | 0.01 | 0.21 |
| | Dawn | 0.00 | 0.05 |
| | Dusk | 0.04 | 0.19 |
| | Night | 0.54 | 0.75 |
| | Darkest night | 0.11 | 0.27 |
| *Red fox* | Day | 0.09 | 0.46 |
| | Dawn | 0.00 | 0.03 |
| | Dusk | 0.03 | 0.20 |
| | Night | 0.34 | 0.70 |
| | Darkest night | 0.07 | 0.36 |
| *Striped skunk* | Day | 0.02 | 0.20 |
| | Dawn | 0.00 | 0.07 |
| | Dusk | 0.02 | 0.32 |
| | Night | 0.39 | 0.77 |
| | Darkest night | 0.08 | 0.49 |
| *Virginia opossum* | Day | 0.00 | 0.18 |
| | Dawn | 0.00 | 0.06 |
| | Dusk | 0.03 | 0.17 |
| | Night | 0.50 | 0.71 |
| | Darkest night | 0.14 | 0.34 |
| *White-tailed deer* | Day | 0.29 | 0.73 |
| | Dawn | 0.02 | 0.09 |
| | Dusk | 0.02 | 0.34 |
| | Night | 0.15 | 0.47 |
| | Darkest night | 0.04 | 0.27 |

Table S4. The mean, standard deviation, and range for each spatial predictor variable collected at three scales across ten U.S. cities.

| | | Avaliable greenspace | | | | Proportion vegetation cover | | | | Proportion impervious cover | | | | Human population density | | | |
|---|---|---|---|---|---|---|---|---|---|---|---|---|---|---|---|---|---|
| | City | Mean | St. Dev. | Min | Max | Mean | St. Dev. | Min | Max | Mean | St. Dev. | Min | Max | Mean | St. Dev. | Min | Max |
| *1500-m buffer* | AUTX | 0.49 | 0.19 | 0.19 | 0.87 | 0.73 | 0.13 | 0.47 | 0.97 | 27.65 | 15.75 | 2.62 | 56.31 | 1888.53 | 1220.69 | 141.68 | 4429.08 |
| | CHIL | 0.23 | 0.21 | 0.00 | 0.81 | 0.62 | 0.23 | 0.15 | 0.98 | 43.01 | 19.50 | 6.78 | 82.30 | 2810.71 | 2679.85 | 0.57 | 10832.15 |
| | DECO | 0.26 | 0.18 | 0.02 | 0.78 | 0.58 | 0.19 | 0.15 | 0.84 | 38.72 | 13.06 | 10.33 | 72.11 | 2152.40 | 1193.33 | 581.71 | 5784.99 |
| | FOCO | 0.27 | 0.11 | 0.13 | 0.60 | 0.66 | 0.12 | 0.40 | 0.86 | 20.58 | 13.46 | 1.37 | 43.01 | 1040.69 | 905.46 | 67.37 | 3284.48 |
| | ICIA | 0.43 | 0.16 | 0.11 | 0.72 | 0.83 | 0.13 | 0.54 | 1.00 | 14.26 | 11.90 | 0.39 | 45.12 | 738.09 | 644.49 | 7.50 | 2402.71 |
| | ININ | 0.31 | 0.18 | 0.03 | 0.70 | 0.80 | 0.15 | 0.40 | 0.99 | 28.28 | 17.79 | 0.75 | 65.97 | 1109.77 | 632.48 | 69.07 | 2491.45 |
| | LBCA | 0.40 | 0.41 | 0.00 | 0.98 | 0.14 | 0.10 | 0.02 | 0.37 | 38.60 | 29.84 | 1.33 | 80.20 | 2056.14 | 1922.56 | 0.71 | 5706.86 |
| | MAKS | 0.50 | 0.24 | 0.13 | 0.99 | 0.90 | 0.08 | 0.65 | 1.00 | 15.97 | 14.23 | 0.16 | 47.33 | 658.39 | 708.23 | 3.54 | 2958.52 |
| | MAWI | 0.20 | 0.14 | 0.01 | 0.52 | 0.71 | 0.21 | 0.11 | 0.94 | 29.41 | 9.76 | 10.00 | 44.29 | 1590.56 | 1039.33 | 319.73 | 5231.02 |
| | WIDE | 0.51 | 0.17 | 0.14 | 0.84 | 0.55 | 0.23 | 0.11 | 0.88 | 22.23 | 15.41 | 0.40 | 59.37 | 1476.80 | 1209.28 | 74.16 | 4891.33 |
| *1000-m buffer* | AUTX | 0.50 | 0.21 | 0.12 | 0.93 | 0.73 | 0.15 | 0.41 | 0.97 | 27.54 | 17.25 | 1.01 | 61.08 | 2115.42 | 1531.26 | 54.46 | 6550.31 |
| | CHIL | 0.24 | 0.23 | 0.00 | 0.89 | 0.63 | 0.24 | 0.14 | 1.00 | 41.82 | 20.45 | 3.07 | 82.59 | 2826.12 | 2935.74 | 0.00 | 13509.83 |
| | DECO | 0.28 | 0.21 | 0.02 | 0.85 | 0.60 | 0.21 | 0.16 | 0.91 | 37.23 | 14.69 | 7.08 | 71.73 | 2293.86 | 1380.39 | 378.01 | 6791.38 |
| | FOCO | 0.29 | 0.12 | 0.10 | 0.64 | 0.64 | 0.15 | 0.37 | 0.91 | 21.55 | 15.01 | 0.85 | 49.01 | 1088.06 | 1007.01 | 47.77 | 3601.73 |
| | ICIA | 0.43 | 0.18 | 0.10 | 0.79 | 0.83 | 0.14 | 0.56 | 1.00 | 14.51 | 13.08 | 0.24 | 45.48 | 844.44 | 739.54 | 8.92 | 2548.92 |
| | ININ | 0.32 | 0.19 | 0.03 | 0.73 | 0.80 | 0.17 | 0.41 | 1.00 | 27.44 | 17.51 | 0.50 | 67.26 | 1187.35 | 725.74 | 97.45 | 2786.48 |
| | LBCA | 0.40 | 0.41 | 0.00 | 0.98 | 0.13 | 0.10 | 0.01 | 0.37 | 38.88 | 30.09 | 1.27 | 83.76 | 2170.22 | 2085.75 | 1.59 | 6358.60 |
| | MAKS | 0.49 | 0.26 | 0.08 | 0.99 | 0.89 | 0.10 | 0.49 | 1.00 | 17.06 | 16.03 | 0.13 | 51.58 | 774.94 | 906.34 | 0.64 | 4597.54 |
| | MAWI | 0.19 | 0.15 | 0.00 | 0.55 | 0.69 | 0.21 | 0.05 | 0.97 | 29.90 | 13.03 | 3.51 | 58.66 | 1761.26 | 1624.73 | 170.06 | 8625.68 |
| | WIDE | 0.53 | 0.18 | 0.14 | 0.88 | 0.54 | 0.26 | 0.02 | 0.91 | 20.66 | 15.72 | 0.32 | 56.22 | 1654.39 | 1450.98 | 38.53 | 5143.05 |
| *500-m buffer* | AUTX | 0.48 | 0.26 | 0.07 | 1.00 | 0.73 | 0.20 | 0.30 | 1.00 | 28.50 | 20.81 | 0.00 | 73.27 | 2683.59 | 2159.21 | 34.39 | 6464.64 |
| | CHIL | 0.30 | 0.27 | 0.00 | 0.96 | 0.67 | 0.25 | 0.02 | 1.00 | 38.10 | 21.83 | 0.44 | 86.09 | 2936.18 | 3087.56 | 0.00 | 13896.12 |
| | DECO | 0.33 | 0.23 | 0.03 | 0.91 | 0.65 | 0.21 | 0.22 | 0.97 | 33.67 | 15.38 | 4.22 | 66.83 | 3002.81 | 1893.46 | 25.48 | 9111.65 |

| | | | | | | | | | | | | | | | | | |
|---|---|---|---|---|---|---|---|---|---|---|---|---|---|---|---|---|---|
| | FOCO | 0.28 | 0.13 | 0.04 | 0.58 | 0.62 | 0.19 | 0.33 | 0.97 | 21.69 | 17.53 | 0.69 | 56.39 | 1300.24 | 1356.83 | 0.00 | 4960.26 |
| | ICIA | 0.48 | 0.22 | 0.10 | 0.92 | 0.86 | 0.13 | 0.56 | 1.00 | 12.21 | 12.77 | 0.01 | 46.70 | 1466.85 | 1312.88 | 25.48 | 5036.69 |
| | ININ | 0.35 | 0.22 | 0.02 | 0.83 | 0.80 | 0.20 | 0.14 | 1.00 | 25.04 | 16.57 | 0.51 | 67.61 | 1465.84 | 946.64 | 20.38 | 3864.77 |
| | LBCA | 0.40 | 0.42 | 0.00 | 0.98 | 0.12 | 0.10 | 0.00 | 0.42 | 39.00 | 30.54 | 1.26 | 82.07 | 3067.47 | 3219.20 | 0.00 | 9502.71 |
| | MAKS | 0.50 | 0.29 | 0.02 | 1.00 | 0.90 | 0.12 | 0.30 | 1.00 | 17.39 | 17.42 | 0.00 | 65.16 | 1068.37 | 1435.39 | 0.00 | 7053.15 |
| | MAWI | 0.20 | 0.21 | 0.00 | 0.67 | 0.70 | 0.26 | 0.00 | 0.99 | 28.79 | 18.81 | 0.00 | 74.13 | 2316.50 | 2664.59 | 0.00 | 13632.44 |
| | WIDE | 0.59 | 0.21 | 0.12 | 0.97 | 0.52 | 0.29 | 0.00 | 0.93 | 16.80 | 14.94 | 0.27 | 57.00 | 2123.78 | 1848.55 | 0.00 | 7105.38 |

Table S5. Name and location of the NOAA weather stations used to calculate daily average temperature for each city.

| City | Station ID | Station Name | Latitude | Longitude |
|---|---|---|---|---|
| Austin, Texas | USW00013904 | AUSTIN BERGSTROM AP | 30.1831 | -97.68 |
| Chicago, Illinois | USW00094846 | CHICAGO OHARE INTL AP | 41.995 | -87.9336 |
| Denver, Colorado | USW00003017 | DENVER INTL AP | 39.8328 | -104.6575 |
| Fort Collins, Colorado | USR0000CRDS | REDSTONE COLORADO | 40.5708 | -105.2264 |
| Iowa City, Iowa | USW00014990 | CEDAR RAPIDS MUNI AP | 41.8833 | -91.7167 |
| Indianapolis, Indiana | USW00093819 | INDIANAPOLIS | 39.7075 | -86.2803 |
| Long Beach, California | USW00023129 | LONG BEACH DAUGHERTY FLD | 33.8117 | -118.1464 |
| Manhattan, Kansas | USW00013996 | TOPEKA MUNI AP | 39.0725 | -95.6261 |
| Madison, Wisconsin | USW00014837 | MADISON DANE RGNL AP | 43.1406 | -89.3453 |
| Wilmington, Delaware | USW00013781 | WILMINGTON NEW CASTLE CO AP | 39.6728 | -75.6008 |
